# Pathogenic bacteria experience pervasive RNA polymerase backtracking during infection

**DOI:** 10.1101/2023.05.12.540596

**Authors:** Kaitlyn R. Browning, Houra Merrikh

## Abstract

Pathogenic bacteria and their eukaryotic hosts are in a constant arms race. Hosts have numerous defense mechanisms at their disposal that not only challenge the bacterial invaders, but have the potential to disrupt molecular transactions along the bacterial chromosome. However, it is unclear how the host impacts association of proteins with the bacterial chromosome at the molecular level during infection. This is partially due to the lack of a method that could detect these events in pathogens while they are within host cells. We developed and optimized a system capable of mapping and measuring levels of bacterial proteins associated with the chromosome while they are actively infecting the host (referred to as PIC-seq). Here, we focused on the dynamics of RNA polymerase (RNAP) movement and association with the chromosome in the pathogenic bacterium *Salmonella enterica* as a model system during infection. Using PIC-seq, we found that RNAP association patterns with the chromosome change during infection genome-wide, including at regions that encode for key virulence genes. Importantly, we found that infection of a host significantly increases RNAP backtracking on the bacterial chromosome. RNAP backtracking is the most common form of disruption to RNAP progress on the chromosome. Interestingly, we found that the resolution of backtracked RNAPs via the anti-backtracking factors GreA and GreB is critical for pathogenesis, revealing a new class of virulence genes. Altogether, our results strongly suggest that infection of a host significantly impacts transcription by disrupting RNAP movement on the chromosome within the bacterial pathogen. The increased backtracking events have important implications not only for efficient transcription, but also for mutation rates as stalled RNAPs increase the levels of mutagenesis.

## Introduction

Antimicrobial resistance is an urgent threat to human health. Bacterial pathogens continue to evolve at a rate that outpaces almost all old and new therapeutics. Resolving this problem requires a general understanding of how hosts and pathogens interact during infection. Numerous studies over the past few decades have demonstrated how different pathogens reach, invade, and proliferate within their target hosts. However, how the host impacts the pathogen during infection at the molecular level, specifically the complexes that function on the bacterial chromosome, has not been thoroughly investigated.

Bacterial pathogens face significant challenges from the moment of initial contact with mammalian hosts. They must successfully survive the harsh host environment and regulate expression of key virulence factors like pathogenicity island-encoded secretion systems, flagella, ion transporters, and stress response genes. For enteric intracellular pathogens, ingested cells must respond to the acidic pH^1^ and the presence of reactive nitrogen species^2^ within the host stomach. They must also overcome nutritional barriers caused by dense microbiota layers^3^ and withstand antimicrobial peptides released by host cells^4^. The bacteria are also heavily bombarded by oxidative stress^5–9^. Although pathogenic bacteria are well equipped to respond to these host defense mechanisms, any number of these stresses during infection could lead to an accumulation of DNA damage, especially oxidative stress. Indeed, studies in *Salmonella enterica*^10,11^, *Staphylococcus aureus*^12^, and *Helicobacter pylori*^13^ demonstrate the critical importance of DNA repair during infection.

DNA damage could have numerous consequences for bacterial cells during infection, including perturbations to DNA replication movement through regions of the chromosome that are packed with stalled RNA polymerases (RNAPs), which increase mutagenesis. Additionally, given that RNAPs are the most abundant proteins associated with the chromosome, DNA damage during infection could have a profound impact on transcription. It is well documented that obstacles on the DNA template, such as damaged template bases or DNA-protein adducts, stall transcription elongation complexes^14–17^. Gene expression programs that respond to the host environment are highly intricate and tightly regulated^18,19^. Thus, any unresolved disruption to RNAP progression during infection could be catastrophic to survival of the pathogen. However, the impact that infection of a host has on RNAP dynamics along the pathogen chromosome has not been investigated. This is partially due to the lack of a precise method capable of detecting changes at the molecular level on the chromosome of pathogens while they are within host cells.

In addition to perturbing transcriptional profiles of pathogenic species during infection, disruptions to RNAP movement could increase mutagenesis. Work from our lab and others’ have shown that RNAP stalling activates transcription-coupled nucleotide repair, which is mutagenic specifically because of endogenous reactive oxygen species^14,20–24^. Therefore, potential disruptions to RNAP movement induced by the host are likely to not only abrogate the transcriptional profile of the pathogen, but also increase the chances of generating hypervirulent strains and facilitating AMR development due to increased mutagenesis.

Here, we investigated how the host impacts RNAP movement on the chromosome of a bacterial pathogen. We used the facultative intracellular pathogen *Salmonella enterica* serovar Typhimurium (*S.* Typhimurium) as our model organism. We developed a novel method, post-infection ChIP-seq (PIC-seq), that can measure RNAP density and changes in RNAP movement on the bacteria chromosome while the pathogen is inside the host cell. We show that RNAP backtracking, the most common form of stalled RNAP complexes (described below), is significantly more prevalent during infection compared to cells grown in broth culture. Furthermore, we find that these disruptions to RNAP progression occur across the genome, including at regions encoding for key *S.* Typhimurium virulence genes. We also show that resolution of backtracking is critical to pathogenesis, highlighting a key role for anti-backtracking factors GreA and GreB in infections. Our results suggest that RNAP movement is substantially impacted by the host, and that the resolution of these disruptions is critical for the survival of pathogens during infection.

## Results

### The development of a method to measure changes in bacterial protein association with the chromosome during infection

We hypothesized that RNAP backtracking is more prevalent during infection genome-wide, and that resolution is critical to pathogenesis. To test this hypothesis, we needed a system that would allow us to map RNAP occupancy on the chromosome of the pathogen while the bacteria still resided within host cells. To our knowledge, such a method did not exist. We developed and optimized a system whereby well-established infection protocols are paired with chromatin immunoprecipitation (ChIP) of bacterial proteins, followed by deep sequencing of the immunoprecipitating DNA (Fig 1A). We refer to this method as post-infection ChIP-seq (PIC-seq). PIC-seq enabled us to determine RNAP dynamics at the molecular level specifically during infection.

**Figure 1.**
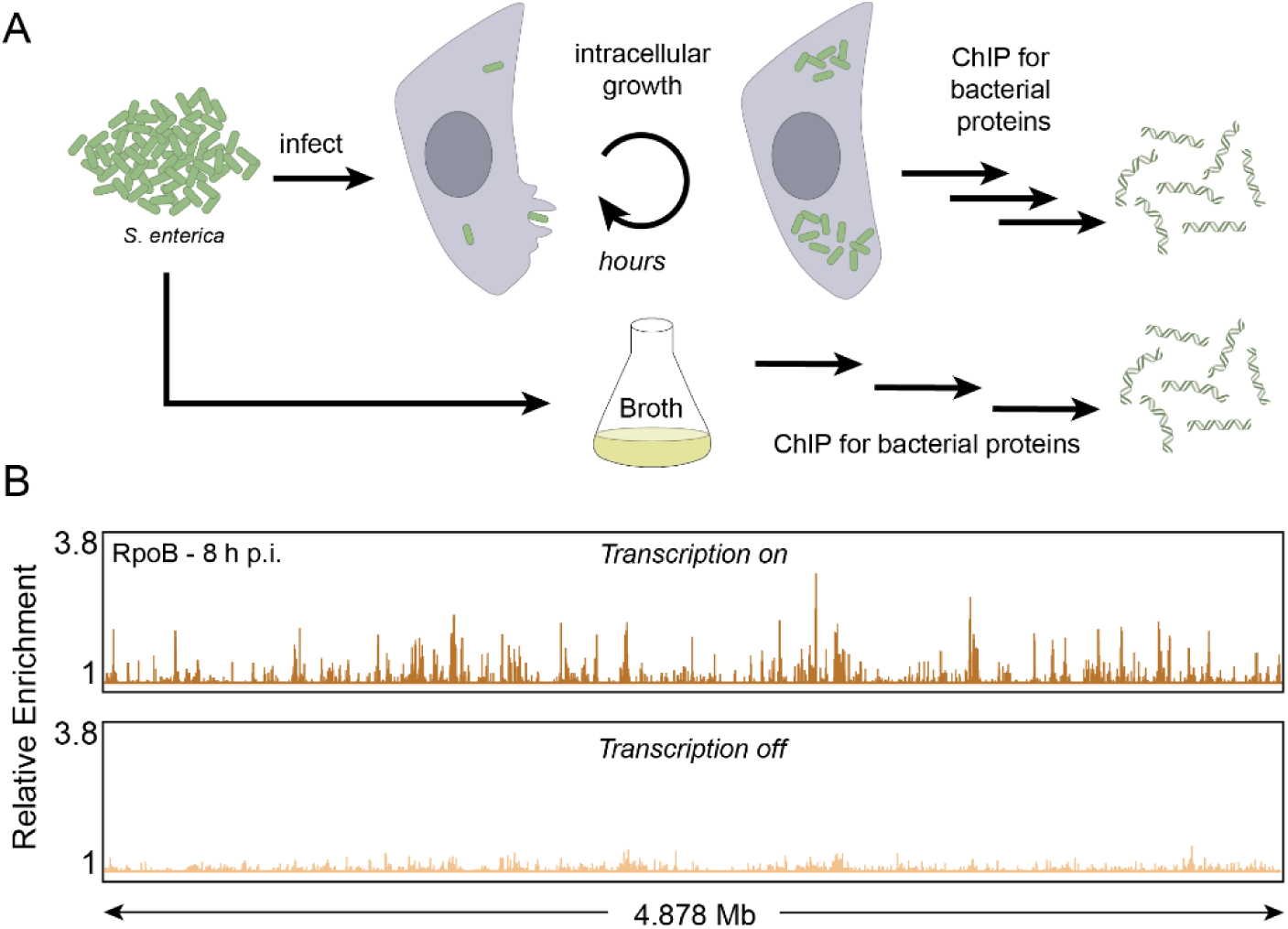
PIC-seq method can be used to measure RNAP occupancy during infection. (A) Schematic of the method to perform chromatin immunoprecipitation (ChIP) followed by deep sequencing from broth culture and from infected cells. Eukaryotic host cells are infected with pathogenic bacteria cells. Once the infection has progressed for a given time, ChIP experiments can be performed. (B) Representative example of *S.* Typhimurium RNAP occupancy 8 h post infection (p.i.) of HeLa cells as determined by PIC-seq of RpoB (the beta subunit of RNAP). Inhibiting transcription with rifampicin during the infection (“transcription off”) abrogates RpoB (RNAP) enrichment genome wide. Relative enrichment is defined as the ratio of IP and total read counts. Total genome size is 4.878 Mb.

We used *Salmonella enterica* serovar Typhimurium (*S.* Typhimurium) to infect HeLa cells based on highly standardized protocols^9,25–27^. At the end of the experiment, bacterial proteins were chemically crosslinked to the DNA while the bacteria still resided within the host cell. Crosslinking the entire infection system in this way ensured that any transient protein interactions within the pathogen that may be unique to infection were not lost during sample processing. Both bacterial and host cells were then lysed and their DNA was fragmented by sonication. RpoB, the β subunit of RNAP, was immunoprecipitated from the mixed lysate using a native antibody. Using Western blot analyses, we determined that the antibody is highly specific to RpoB and does not cross react with proteins specific to HeLa cells (S1 Fig). The associated DNA was then extracted and deep sequenced. Any contaminating DNA from the host is naturally excluded from further analysis, as it does not map to the *S.* Typhimurium genome. On average, 12.4% of total reads mapped to the *S.* Typhimurium genome in our experiments, providing an average of approximately 80x coverage (approximately 2.6 million aligned reads per sample). Though we used this system to immunoprecipitate RNAP from *Salmonella*, PIC-seq can, in principle, be adapted to examine any protein of interest with a specific antibody in any host-pathogen combination.

We measured RNAP association with the pathogen genome with PIC-seq. In parallel, as a control to determine if our signal was specific to the bacterial pathogen, we treated cells with the antibiotic rifampicin, which is a transcription inhibitor, during the infection. Upon rifampicin treatment, the signal for RNAP occupancy was abrogated genome-wide, demonstrating the specificity of PIC-seq (Fig 1B).

### RNAP association with virulence genes can be specifically detected during infection

To further confirm that our method can differentiate between cells grown in broth culture and infection, we analyzed the levels of RNAP occupancy with regions encoding virulence genes (Fig 2A, S2A Fig). As expected, RNAP was significantly enriched at regions of the chromosome containing multiple, successive virulence genes (known as *Salmonella* pathogenicity islands, “SPIs”) during infection (Fig 2B). Importantly, no significant RNAP signal was found at these genes in cells grown in broth (Fig 2C). Similar trends were observed at a trio of virulence genes, *pipB2*, *virK*, and *mig-14*, where there is no RNAP enrichment in cells grown in broth, but clear enrichment during infection. Unsurprisingly, almost all genes with significant RNAP enrichment during infection (but not in broth culture) have a role (confirmed or putative) in virulence (S1 Table). These results are consistent with previous RNA-seq datasets of *S.* Typhimurium during infection^9,28,29^, as well as the well-defined roles of SPI-1 and SPI-2 proteins in facilitating host cell invasion and promoting intracellular survival, respectively^30^. In contrast, we observed similar levels of RNAP at housekeeping gene regions, such as at rDNA genes, in both conditions (S2B Fig, S2 Table). Our results demonstrate that PIC-seq can specifically map and measure the levels of RNAP occupancy during infection. Furthermore, the high RNAP occupancy at virulence genes we observed further supports prior work and the conclusion that these genes are vital during infection.

**Figure 2.**
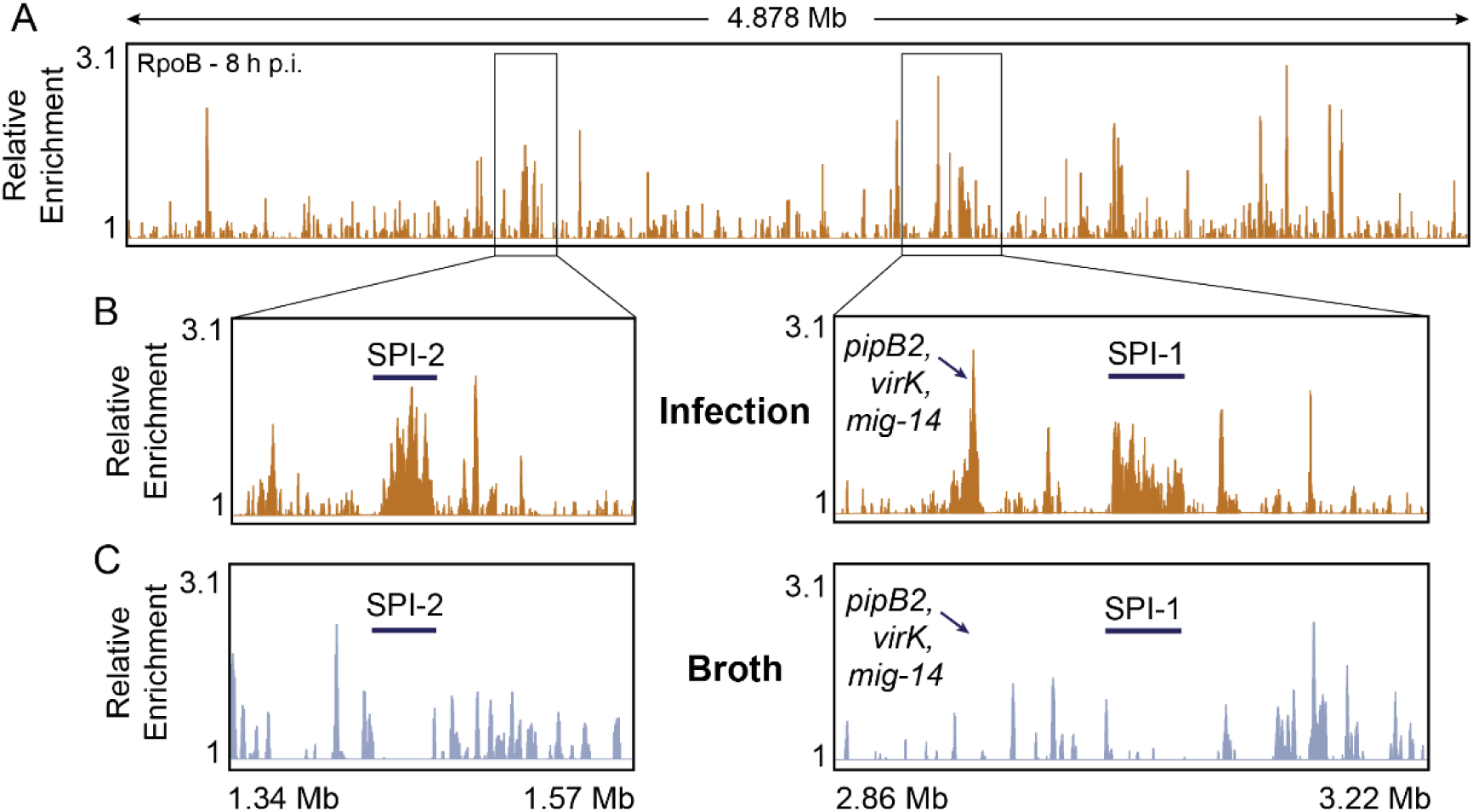
Specific detection of RNAP occupancy at *S.* Typhimurium virulence genes. (A) Representative example of *S.* Typhimurium RNAP occupancy 8 h post infection (h p.i.) of HeLa cells as determined by ChIP-seq of RpoB. (B) Reads mapped to the *S.* Typhimurium genome display areas of high RNAP enrichment during infection, notably within *Salmonella* pathogenicity islands (SPIs) and at a trio of virulence genes (*pipB2*, *virK*, and *mig-14*). (C) No significant RNAP signal was found at these genes in cells grown in broth culture. Relative enrichment is defined as the ratio of IP and total read counts.

### RNAP backtracking increases significantly during infection

We hypothesized that RNAP backtracking is more prevalent during infection genome wide. We postulated this based on the fact that the host can damage DNA (such as through oxidative stress), potentially inhibiting RNAP movement. The most common type of RNAP stalling is backtracking, which is a stable state of the complex^31^. When backtracked, the active site of RNAP becomes disengaged, and the 3’ end of the nascent RNA is extruded through the secondary channel of the enzyme^32–34^. In *S.* Typhimurium, the anti-backtracking factors GreA and GreB rescue backtracked RNAPs by stimulating the intrinsic endonucleolytic activity of RNAP to restore the active site^35,36^.

To test the hypothesis that there is increased RNAP backtracking on the genome of the pathogen during infection, we constructed strains that lack GreA and GreB^37^. If our hypothesis is correct, then cells lacking the Gre factors would have increased levels of stalled RNAPs during infection compared to growth in broth cultures. To detect this, we measured the differences in RNAP occupancy levels across the genome in the presence (wild-type, WT) and absence (Δ*greA* Δ*greB*) of these anti-backtracking factors. Since both Gre factors are directly and specifically involved in the resolution of backtracked RNAPs^35,36^, we deduced that sites where RNAP occupancy changes in the absence of Gre factors are sites where backtracking is occurring. The prevalence of backtracking can then be measured by determining how these RNAP occupancy differences in the presence and absence of Gre factors change in cells grown in broth versus during infection.

We measured RNAP occupancy in the presence and absence of Gre factors in cells grown in broth and at 1 or 8 h p.i. using PIC-seq. Indeed, deletion of both Gre factors led to changes in RNAP occupancy genome-wide (Fig 3A, S3A-B Fig). We then identified genes that exhibited differential RNAP occupancy in the presence and absence of Gre factors in each condition. However, because cells grown in these conditions have widely different gene expression profiles, we could not directly compare backtracking in the same gene across each condition. In other words, we could not compare backtracking at a gene that is transcriptionally active in one condition but not in the other. Therefore, we categorized the genes by transcription level, which was determined for each gene by normalizing the number of reads mapping to that gene in the IP sample versus the total sample in the WT strain. Genes were categorized independently for each condition. This resulted in 631 top transcribed genes in broth, 645 at 1 h p.i., and 590 at 8 h p.i. (S3 Table, S3C Fig). When comparing the 283 top transcribed genes that are shared between all three conditions, it is clear that RNAP occupancy differs in the presence and absence of Gre factors, not only in each independent condition, but also across conditions (Fig 3B).

**Figure 3.**
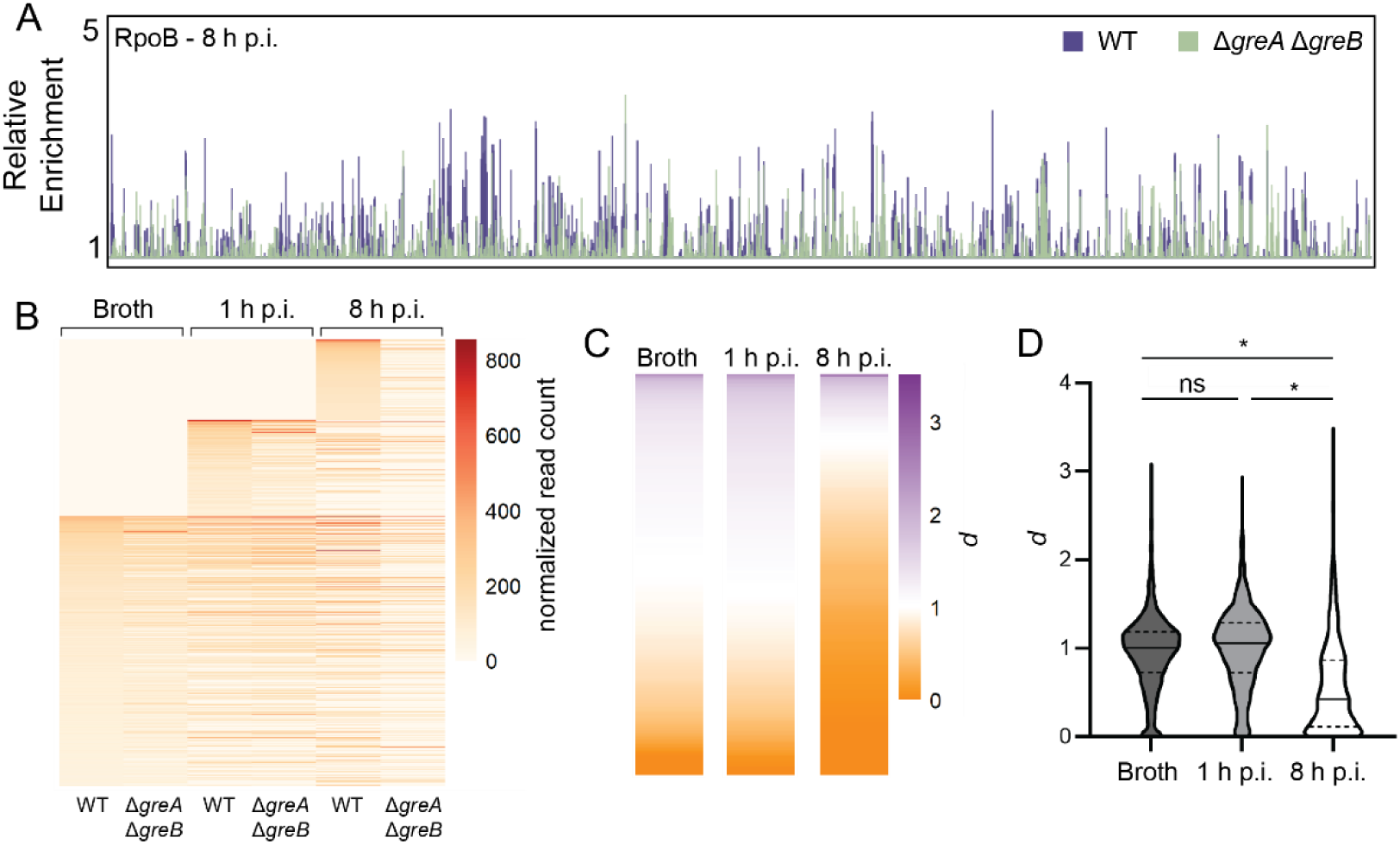
RNAP occupancy decreases in the absence of Gre factors. (A) Representative RNAP occupancy profile for wild-type (WT) cells or cells lacking Gre factors (Δ*greA* Δ*greB*) 8 h post infection (p.i.) of HeLa cells as determined by ChIP-seq of RpoB. Relative enrichment is defined as the ratio of IP and total read counts. (B) RNAP occupancy changes visualized as a heat map, where every horizontal line represents the normalized read count for the same gene across each condition. All top transcribed genes for each condition are plotted (1043 genes total, see S3 Table). Each value represents the average of three biological replicates. Zeros are plotted for genes that do not arise in one condition but do arise in one or both of the other conditions. (C) Ratio *d* of normalized read count in the absence of Gre factors versus in the presence of Gre factors (WT) for top-transcribed genes in each condition averaged across three biological replicates. (D) Quantification of (C) showing ratio *d* calculated for every top-transcribed gene, where the median is shown by a solid line and quartiles by dashed lines. \**p*<0.0001, ns: not significant, one-way ANOVA.

We found that RNAP occupancy generally decreases in the absence of Gre factors. This is consistent with the activity of previously described accessory factors, such as helicases, that can remove stalled RNAPs from DNA^16,21, 38–42^. Backtracked RNAPs that are not rescued by the Gre factors are expected to eventually become a target for removal by such helicases and other transcription terminators, leading to the decrease in RNAP occupancy that we observe.

We performed a more quantitative analysis to get an accurate picture of RNAP occupancy changes for cells grown during infection versus growth in broth culture. We defined a ratio, *d*, as the normalized read counts mapping to a gene in the backtracking-prone state (Δ*greA* Δ*greB*) versus those in WT. An equivalent ratio (*d* ≈ 1) indicates that there is little difference in RNAP occupancy independent of the presence of the Gre factors. We interpreted these regions to be those that are not experiencing backtracking. A ratio that deviates from one (*d* < 1 or *d* > 1) indicates that there is a difference in RNAP occupancy at the gene that is dependent on the Gre factors, suggesting that these factors are playing an important role in maintaining RNAP progression at that gene. We interpreted these regions to be those that are experiencing significant RNAP backtracking. This ratio was calculated for every top-transcribed gene in each condition (Fig 3C, S3D Fig). For cells grown in broth and at 1 h p.i., the average ratio of RNAP occupancy in the absence of Gre factors versus in the presence is equivalent (Table 1, Fig 3D), suggesting that, on average, cells do not depend on Gre factors to maintain proper RNAP occupancy under these conditions. In contrast, cells at 8 h p.i. do rely on the Gre factors to maintain RNAP occupancy (Table 1, Fig 3D), suggesting that backtracking becomes pervasive as the infection progresses.

**Table 1.** Gre factor-dependent RNAP occupancy changes in broth versus infection.

| Transcription level | $d_{broth}$ | $d_{1 \text{ h p.i.}}$ | $d_{8 \text{ h p.i.}}$ | $d'$<br>1 h p.i. vs. broth | $d'$<br>8 h p.i. vs. broth |
| --- | --- | --- | --- | --- | --- |
| ++ | 0.96 | 1.01 | 0.56 | 1.06 | 0.59 |
| +++ | 0.91 | 0.99 | 0.52 | 1.09 | 0.57 |
| ++++ | 0.91 | 0.94 | 0.48 | 1.03 | 0.53 |
| All genes | 0.94 | 1.00 | 0.55 | 1.06 | 0.59 |

Ratio *d* was calculated as the normalized read counts mapping to a gene in the backtracking-prone state (Δ*greA* Δ*greB*) versus in WT for each condition and each transcription level (++, +++, ++++). Ratio *d’* comparing ratio *d* for 1 or 8 h p.i. to broth for the groups of genes with equivalent transcription levels.

We determined the extent to which backtracking is more prevalent during infection than broth. We accomplished this by quantifying the average differences in RNAP occupancy changes in broth versus infection. We defined a new ratio, *d’*, as the ratio of *d* for infection versus *d* for broth, *d’* = *d_inf_ / d_broth_.* Backtracking early in infection, at 1 h p.i., is not more prevalent than in broth, but it is more prevalent at 8 h p.i., *d’* = 0.59 (Table 1).

### Backtracking occurs within key genes necessary for *Salmonella* pathogenesis

Having established that backtracking is more prevalent during infection genome-wide, specifically at later time points, we endeavored to identify those regions where backtracking occurs during infection. Thus, we looked at the genes where the greatest changes in RNAP occupancy in the absence of Gre factors occur. These were identified by plotting RNAP occupancy as heat maps and using hierarchical clustering (Fig 4A, S4 Fig, S4 Table).

**Figure 4.**
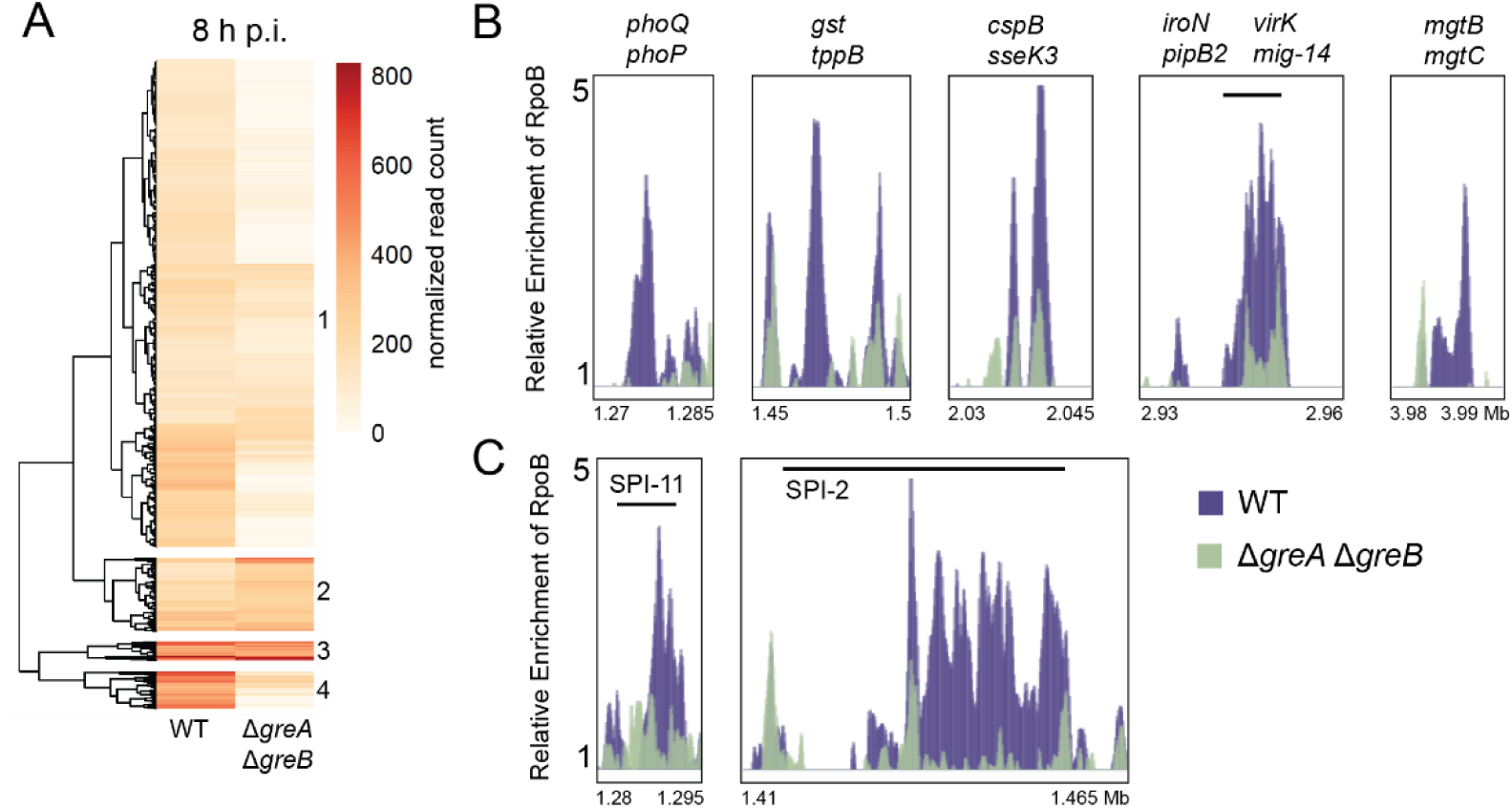
Backtracking is prevalent at key virulence genes during infection. (A) RNAP occupancy changes at 8 h post infection (p.i.) in the absence of Gre factors visualized as a heat map. Each value represented is the average of three biological replicates. Hierarchical clustering (numbered 1-4) was performed using the pheatmap function in RStudio (see S4 Table). (B) Key genes related to virulence in cluster 4 in (A) exhibit significant differences in RNAP occupancy (*p* < 0.05 for every gene listed, unpaired t test). (C) Genes belonging to SPIs that fall into cluster 1 in (A) and exhibit significant differences in RNAP occupancy.

As backtracking is minimal throughout all clusters in broth and at 1 h p.i. (Fig 3, S4 Fig), we focused on characterizing gene functions for each cluster at 8 h p.i. (Fig 4A). Genes at 8 h p.i. fall into four clusters: cluster one, 466 genes; cluster two, 70 genes; cluster three, 18 genes; and cluster four, 36 genes (S4 Table). In clusters four and one, RNAP occupancy is generally higher in the presence of Gre factors (WT) than in the absence (Δ*greA* Δ*greB*). Genes in cluster four exhibit dramatic differences in RNAP occupancy depending on the presence of Gre factors, with an average *d* of 0.23. There was not enough statistical power to perform gene ontology (GO) functional enrichment analysis for genes in cluster four. However, manual annotation of these gene functions reveals that out of 36 genes, at least 15 have been previously implicated in *S.* Typhimurium virulence (Fig 4B, S4 Table). Although RNAP occupancy does depend on the presence of Gre factors in cluster one (*d* = 0.43), these genes have lower RNAP occupancy levels in both genotypes (Fig 4A). GO functional enrichment analysis revealed that genes in cluster one are enriched in functions related to cellular and macromolecular localization, membrane biogenesis, regulation of metabolic processes, secretion, and translation. This cluster also consists of numerous SPI genes that exhibit large differences in RNAP occupancy when Gre factors are absent (Fig 4C, S4 Table). Of the 25 top transcribed SPI-2 genes in this dataset, 22 fall within cluster one. Similarly, all five of the top transcribed SPI-11 genes in this dataset fall within cluster one. Efficient RNAP progression at these genes would be critical to ensuring cell survival during infection, as these genes are required throughout *S.* Typhimurium infection.

Genes in cluster two exhibit an average *d* ratio of 1.41, or greater RNAP occupancy in the absence than in the presence of Gre factors. Genes in cluster two are enriched in functions related to cellular nitrogen compound biosynthetic processes (GO analysis). Cluster three includes genes where RNAP occupancy is high and does not change in the absence of Gre factors (*d* = 0.99). Manual annotation of genes in cluster three reveal functions related to essential processes such as translation, carbon metabolism, and cell wall maintenance (S4 Table). Overall, our results suggest that backtracking occurs at key virulence genes during infection, and that the Gre factors are critical for RNAP progression at these regions.

We found that the prevalence of backtracking was not uniform genome-wide. We reasoned that longer genes are more likely to accumulate DNA damage and therefore more likely to accumulate backtracked RNAPs. Accordingly, we found that the average length of genes in cluster four, which exhibited the greatest differences in RNAP occupancy in a Gre factor-dependent manner, was significantly longer than the average length of all top-transcribed genes in this dataset (S5 Fig).

### Resolution of backtracking is critical to pathogenesis

Knowing that backtracking is prevalent at key virulence genes during infection, we tested whether disruption to RNAP progression is problematic for pathogenesis in general. To elucidate the impact of prevalent backtracking on *S.* Typhimurium pathogenicity, we infected HeLa cells with backtracking-prone *S.* Typhimurium lacking one or both Gre factors. We measured the number of viable intracellular *S.* Typhimurium from the infection over time using a standard gentamicin protection assay (Fig 5). Deletion of *greB* alone did not significantly affect the growth of *S.* Typhimurium inside HeLa cells over the course of the infection, while deletion of *greA* led to a modest, yet statistically significant, loss in intracellular growth. However, deletion of both Gre factors significantly abrogated intracellular growth of *S.* Typhimurium throughout the duration of the infection (Fig 5A). This effect is specific to infection, as there were no significant differences in growth of these strains compared to wild type when grown in broth (Fig 5B). The infection-specific effect of depleting cells of both Gre factors suggests that pervasive backtracking during infection disrupts key virulence gene expression, and resolution is critical to intracellular survival.

**Figure 5.**
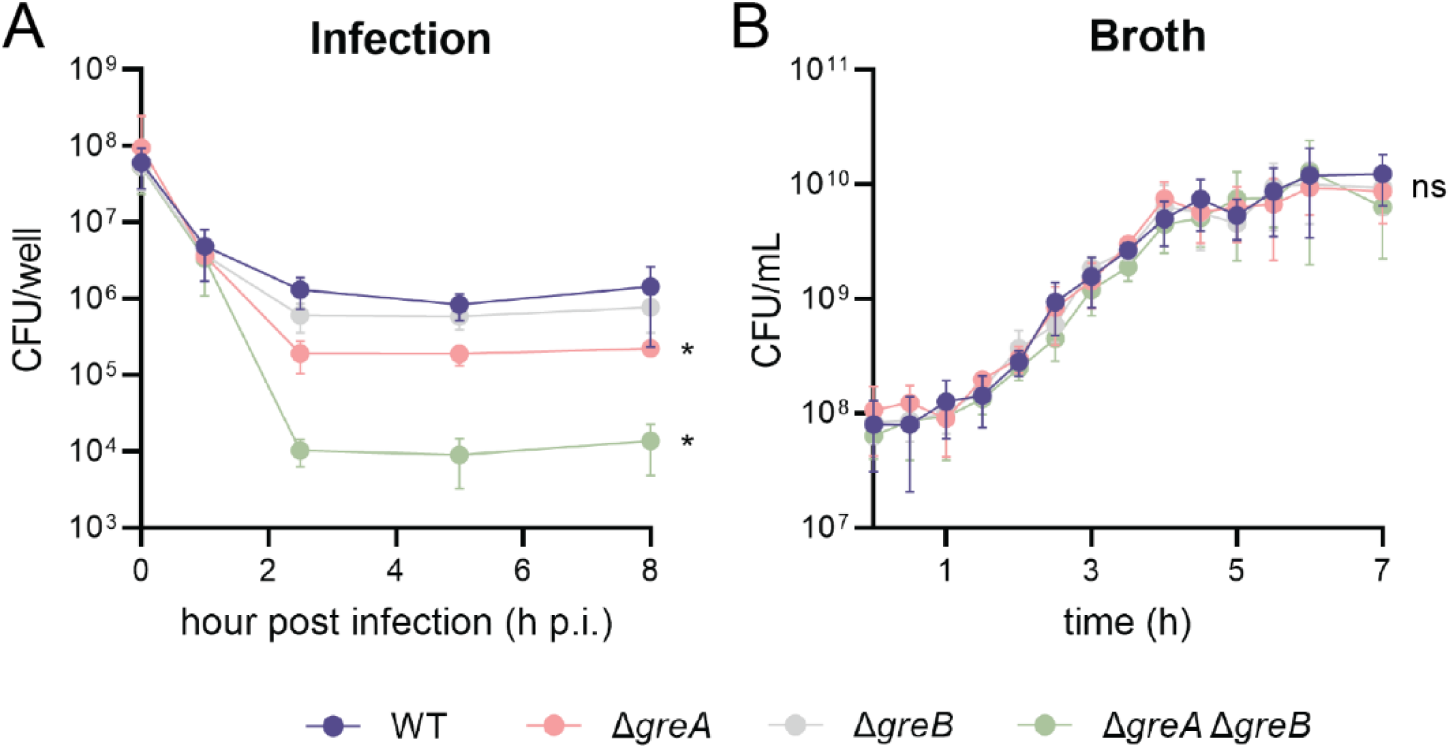
Resolution of backtracking during infection is key to pathogenesis. Average number of CFUs of backtracking-prone cells (A) isolated during infection or (B) grown in broth culture. Error bars indicate standard deviation of at least six biological replicates. \**p*<0.005, ns: not significant, one-way ANOVA with Bonferroni correction.

## Discussion

During infection, bacterial pathogens must survive numerous host defense mechanisms that are not only damaging to the cell itself, but can also lead to DNA damage capable of stalling RNAP^14–17^. Pathogens must quickly sense and respond to the host environment, and any disruption to these virulence gene programs at the transcriptional level would be significant. Our work suggests that infection of a host significantly impacts bacterial transcription, causing pervasive backtracking that must be resolved to ensure successful pathogenesis.

Our data show that disruptions to RNAP movement increase as the infection progresses. As evidenced by the lack of difference in RNAP occupancy in the presence and absence of Gre factors, it appears that after 1 h of infection, RNAPs are not yet perturbed by interactions with the host. This is further emphasized by growth of cells at 1 h p.i.: cells at this time point do not require Gre factors to survive during infection. It is possible, however, that RNAP stalling at 1 h p.i. is underestimated in our experiments due to detection thresholds of PIC-seq as a population assay and the heterogeneity of the cell population at this early point post infection^43^. Cells at 8 h p.i., however, have endured stress from the host environment longer than those at 1 h p.i., potentially increasing the likelihood of DNA damage accumulation that will stall RNAPs and lead to backtracking. Cells at this later time point exhibit large differences in RNAP occupancy and, based on infection experiments in the presence and absence of Gre factors, require backtracking resolution for intracellular survival.

In addition to perturbing transcription, an increase in the prevalence of backtracking during infection could have implications for DNA replication. Transcription is an obstacle to DNA replication due to the lack of spatiotemporal separation between the two processes. These conflicts pose significant threats to genome stability regardless of directionality. Interestingly, it has been demonstrated that co-directional conflicts between a backtracked RNAP and an oncoming replication fork lead to double-stranded breaks (DSBs) in the DNA^37^. Our data show that *S.* Typhimurium cells lacking previously described key conflict resolution factors, such as UvrD or Rep helicases^44^, exhibit significant growth defects during infection of HeLa cells (S6 Fig). However, further work is required to determine the extent to which replication-transcription conflicts occur in pathogens during infection. Whether increased backtracking during infection exacerbates conflicts, thus compromising genome integrity, is untested. Previous work has shown that cells that are prone to backtracking (lacking either Gre factor) accumulate mutations faster^37^ and have higher rates of recombination than cells with both Gre factors^45^.

Previous literature shows that stalled RNAPs trigger downstream DNA repair pathways that can be mutagenic^14,20–24^. Although pathogenic cells do have the Gre factors and will thus experience less backtracking under regular conditions, the fact that these RNAP stalling events are much more prevalent during infection inherently increases the chances of mutagenic events occurring. Therefore, we postulate that, though the host is fighting the bacteria with DNA damaging mechanisms such as oxidative stress, they are also enabling bacteria to attain mutations that could lead to adaptive evolution. This idea is outside the scope of this work; however, it is certainly interesting to investigate in future studies as this dichotomy could accelerate the development of AMR and/or hypervirulence.

Our results also highlight an underappreciated role for Gre factors during infection and suggest the existence of a new class of so-called virulence genes in bacterial pathogens. This class of proteins promote survival and guarantee proper gene expression during infection by regulating and modulating RNAP through transcription initiation, elongation, and termination. Consistent with our findings, in *S.* Typhimurium, Gre factors have been associated with promoting intracellular fitness during oxidative stress by facilitating transcriptional fidelity and elongation of central metabolic genes^46^. They have also been demonstrated to prevent backtracking within one gene, *hilD*, which encodes a master regulator of SPI genes to promote pathogenesis^47^, and to facilitate biofilm formation through mediating transcription elongation at key effector genes^48^. Similar impact of Gre factors on virulence phenotypes have also been shown in other, highly divergent bacterial species which highlights the conservation of this phenomenon^49–51^.

Precise regulation of gene expression during infection is critical to the survival of bacterial pathogens. Our data strongly suggest that DNA damage to the pathogen resulting from host defense mechanisms threatens this precision. Disruptions to RNAP progression must be resolved. Altogether, our results support the model that backtracking is prevalent during infection and that Gre factors are required to ensure proper gene regulation and survival during infection.

## Methods

### Bacterial strains and growth conditions

*Salmonella enterica* serovar Typhimurium SL1344 was the wild-type (WT) strain used in these studies. Derivative mutant strains are listed in S5 Table. Bacteria were grown on LB-Lennox agar plates at 37°C with the following antibiotics when appropriate: 25 mg/mL chloramphenicol or 50 mg/mL kanamycin. Single colonies were used to inoculate liquid cultures in LB-Lennox and grown at 37°C with aeration (260 rpm).

### Construction of chromosomal deletion mutants

The bacteriophage λ Red recombination system was used for construction of chromosomal deletion mutants^52,53^. The plasmid pSIM27 was transformed into wild-type SL1344 for expression of the Red recombinase system. Kanamycin and chloramphenicol resistance cassettes were amplified with Phusion High-Fidelity polymerase (Thermo) from T-SACK gDNA^54^ using primers that contained 40 nucleotides homologous to the regions flanking the *greA* or *greB* (including the ATG start codon, S5 Table). Amplicons were electroporated separately into SL1344 and cells were grown on selective LB agar to yield single knockout strains Δ*greA* (*greA::*Kan, HM4527) and ΔgreB (*greB::*Cat, HM4525). To make the double knockout strain Δ*greA* Δ*greB* (HM4529), the *greA* gene was replaced with the kanamycin resistance cassette in the ΔgreB strain as above. Every deletion was confirmed by Sanger sequencing.

### Mammalian cell culture

HeLa cells (ATCC) were grown in high glucose Dulbecco’s minimal essential medium (DMEM, Gibco – 11995065) supplemented with 10% heat inactivated fetal bovine serum (FBS, R&D Systems), 4 mM L-glutamine, and 1X Penicillin/Streptomycin (Gibco – 15140122). Antibiotic-free media of the same formulation was used for bacterial infections. Cells were maintained at 37°C with 5% CO_2_ and passaged following ATCC guidelines.

### Seeding and bacterial invasion of mammalian cells

HeLa cells at low passage number were seeded at the following densities 16-18 hours before infection: 1.5 × 10^5^ cells per well (24-well dish) or 1.14 × 10^7^ (15-cm plate). Immediately prior to infection, HeLa cells were washed with 1X PBS (Gibco), and antibiotic-free media was added to the cells. Cultures of *S.* Typhimurium were inoculated in LB from single colonies and grown overnight at 37°C while shaking. The next day, the precultures were diluted back to OD_600_ = 0.05 in LB and grown at 37°C while shaking until OD_600_ reached 0.6 (approximately 2 h). Bacteria were collected by centrifugation, washed in 1X PBS, resuspended in antibiotic-free media, and used immediately to infect HeLa cells at an MOI of ∼100:1. Bacteria were allowed to invade for 1 h at 37°C with 5% CO_2_. Bacteria were removed after invasion, and HeLa cells were washed once with 1X PBS and fresh antibiotic-free media was added. Thirty minutes later (1.5 h post infection [p.i.]), gentamicin was added to a final concentration of 50 ug/mL for the duration of the experiment. Infected HeLa cells were maintained at 37°C with 5% CO_2_ until times indicated below.

### Gentamicin protection assays

HeLa cells were seeded in 24-well dishes and infected with bacteria as described above. Extracellular growth of *S.* Typhimurium was inhibited by gentamicin as described above. At indicated time points, infected HeLa cells were washed once with 1X PBS and lysed with ice-cold 1% Triton X-100 in H_2_O. Viable bacteria were enumerated by plating on LB agar and grown at 37°C overnight. A one-way ANOVA was performed at 2.5, 5 and 8 h p.i. timepoints with Bonferroni correction to determine statistical significance.

### Chromatin immunoprecipitation from infection

HeLa cells were seeded in 15-cm plates and infected with bacteria as described above. For treatment with rifampicin, 100 mg/mL was added to the media ten minutes prior to each time point. At indicated time points, infected HeLa cells were washed once with 1X PBS and crosslinked in 1X PBS + 1% methanol-free formaldehyde (Thermo) in H_2_O for 10 min at room temperature and subsequently quenched with 0.5 M glycine. Crosslinked infected HeLa cells were washed twice with 1X PBS, then dislodged in 1X PBS by scraping. Cells from two 15-cm plates were combined for each replicate and collected by centrifugation. Cell pellets were stored at −80°C for future processing. Pellets were thawed on ice and resuspended in 2.5 mL ice-cold NPT lysis buffer^55^ [(50 mM Tris–HCl pH 7.5, 150 mM NaCl, 5 mM EDTA, 0.5 % NP-40, 0.1 % Triton X-100, complete protease inhibitor cocktail [Roche] added fresh) for 10 min on ice. Lysozyme was added to 10 mg/mL and the lysate was incubated at 37°C for 30 min. Lysates were sonicated for 5 cycles of 30 s on/off (2.5 min total sonication time) at 4°C in a Bioruptor Plus sonication system (Diagenode) and pelleted by centrifugation at 8000 RPM for 15 min at 4°C. A 40 μL aliquot was taken from the lysate supernatant as the input control. For the immunoprecipitation, 6 μL RpoB monoclonal antibody (8RB13, Thermo) was added to the lysate supernatant and rotated overnight at 4°C. The next day, 90 μLof a 50% protein A Sepharose bead slurry (GE) was added, and IPs were incubated for 1 h at room temperature with gentle rotation. Beads were pelleted by centrifugation at 2000 RPM for 1 min. The supernatant was discarded, and the beads were washed six times for 3 min each in wash buffer (50 mM Tris-HCl pH 7.0, 150 mM NaCl, 5 mM EDTA, 1% Triton X-100), followed by one wash with TE pH 8.0. The elution was carried out at 65°C for 10 min in 200 uL elution buffer I (50 mM Tris pH 8.0, 10 mM EDTA, 1% SDS). Beads were pelleted at 5000 RPM for 1 min and the supernatant saved. The beads were washed with 150 uL elution buffer II (10 mM Tris-HCl pH 8.0, 1 mM EDTA, 0.67% SDS) and pelleted at 7000 RPM for 1 min. The second supernatant was combined with the first eluate. The combined eluates and the accompanying input controls were de-crosslinked overnight by incubation at 65°C. The following day, the eluates and input controls were treated with proteinase K (0.4 mg/mL) at 37°C for 2 h. Sodium acetate was added and the DNA purified by phenol:chloroform:isoamyl alcohol extraction. The DNA was precipitated in 100% ethanol at −20°C and pelleted at 14,000 RPM for 15 min at 4°C, before being resuspended in resuspension buffer (10 mM Tris-HCl pH 8.0, 1 mM EDTA).

### Chromatin immunoprecipitation from broth

Cultures of *S.* Typhimurium were inoculated in LB from single colonies and grown overnight at 37°C while shaking. The next day, the precultures were diluted back to OD_600_ = 0.05 in LB and grown at 37°C while shaking until OD_600_ reached 0.6 (approximately 2 h). Bacteria were crosslinked with 1% formaldehyde for 20 min at room temperature before being quenched with 0.5 M glycine. Cells were collected by centrifugation and washed once in cold 1X PBS. Cell pellets were resuspended in 1.5 mL Solution A (10 mM Tris–HCl pH 8.0, 20% w/v sucrose, 50 mM NaCl, 10 mM EDTA, 10 mg/ml lysozyme, 1 mM AEBSF) and incubated at 37° C for 30 min. After incubation, 1.5 mL of 2X IP buffer (100 mM Tris pH 7.0, 10 mM EDTA, 2% Triton X-100, 300 mM NaCl and 1 mM AEBSF, added fresh) was added and lysates were chilled on ice for 30 min. Lysates were sonicated four times for 10 s (40 s total sonication time) at 30% amplitude and pelleted by centrifugation at 8000 RPM for 15 min at 4°C. A 40 uL aliquot was taken from the lysate supernatant as the input control. For the immunoprecipitation, 2 uL of RpoB monoclonal antibody (clone 8RB13, Thermo) was added to 1 mL of the lysate supernatant and rotated overnight at 4°C. The next day, 30 uL of 50% protein A Sepharose bead slurry (GE) was added, and IPs were incubated for 1 h at room temperature with gentle rotation. Beads were pelleted by centrifugation at 2000 RPM for 1 min. The supernatant was discarded, and the beads were washed six times for 3 min each in wash buffer (50 mM Tris-HCl pH 7.0, 150 mM NaCl, 5 mM EDTA, 1% Triton X-100), followed by one wash with TE pH 8.0. The elution was carried out at 65°C for 10 min in 100 uL elution buffer I (50 mM Tris pH 8.0, 10 mM EDTA, 1% SDS). Beads were pelleted at 5000 RPM for 1 min and the supernatant saved. The beads were washed with 150 uL elution buffer II (10 mM Tris-HCl pH 8.0, 1 mM EDTA, 0.67% SDS) and pelleted at 7000 RPM for 1 min. The second supernatant was combined with the first elution. The combined eluates and the accompanying input controls were de-crosslinked overnight by incubation at 65°C. The following day, the eluates and input controls were treated with proteinase K (0.4 mg/mL) at 37°C for 2 hr. Sodium acetate was added and the DNA purified by phenol:chloroform:isoamyl alcohol extraction. The DNA was precipitated in 100% ethanol at −20°C for 1 hr and pelleted at 14,000 RPM for 15 min at 4°C, before being resuspended in resuspension buffer (10 mM Tris-HCl pH 8.0, 1 mM EDTA).

### Deep sequencing and data processing

Sequencing libraries were prepared using a Nextera XT DNA Library Preparation kit (Illumina). Libraries were deep sequenced by the Vanderbilt Technology for Advanced Genomics (VANTAGE) sequencing core on an Illumina NovaSeq platform, resulting in approximately 24M x 150 bp paired-end reads per sample. Raw reads were trimmed (Trimmomatic v0.39^56^) then mapped to the *S. enterica* serovar Typhimurium strain SL1344 genome (GenBank: FQ312003.1) using Bowtie2 v2.2.5^57^. Both PCR and optical duplicates were removed using Picard v1.3 (Broad Institute) and bam files were sorted and indexed using SAMtools v1.13^58^.

The number of reads mapping to each gene in both the IP and input samples was quantified using featureCounts v2.0.3^59^ and then normalized to total mapped reads. For each gene, the normalized number of reads in the input sample was subtracted from the normalized number of reads in the IP sample and averaged across three biological replicates to calculate normalized read count. For any gene where this calculation resulted in a negative number, the normalized read count was redefined as 0 for that gene. At this point, rDNA genes were excluded from further analysis. Genes were then categorized into five groups based on transcription level using K-means clustering for each individual condition. Transcription level was defined by the normalized read count in the wild-type sample for each condition. The three groups containing the most transcribed genes for each condition were used for further analysis. Ratio *d* was calculated for these genes by dividing the normalized read count for the Δ*greA* Δ*greB* strain by the normalized read count in the wild-type strain. Ratio *d’* was calculated by dividing *d* from the 1 h p.i. or 8 h p.i. condition by the *d* from the broth condition. Heatmaps were plotted and hierarchical clustering was performed using the pheatmap package in RStudio. Gene ontology functional enrichment analysis was performed using the PANTHER v.14 pipeline^60^ through The Gene Ontology Resource^61,62^.

For enrichment analysis, peaks were called from processed bam files using macs2 v.2.2.7.1^63^ and assigned to genome features using the “closest” tool from the BEDTools suite v.2.30.0^64^.

Bedgraph files were prepared from the processed bam files using the bamCompare tool from the deepTools suite v3.5.1^65^ and visualized on the Integrated Genomics Viewer (igv) platform v2.12.1^66^.

All statistical tests were performed in GraphPad Prism v9.

## Acknowledgments

We would like to thank all past and present members of the Merrikh lab for their suggestions and advice on this project, especially Kevin Lang and Anna Johnson. We are also indebted to the Cortez and Hodges labs for providing their equipment and expertise throughout this project. This project was funded by T32ES007028 to K.R.B. NIH National Institute grant AI127422 to H.M.

## Contributions

H.M. and K.R.B. designed the experiments, K.R.B. performed the experiments, both H.M. and K.R.B. analyzed the data and wrote the paper.

## Supplementary Information

**S1 Figure.**
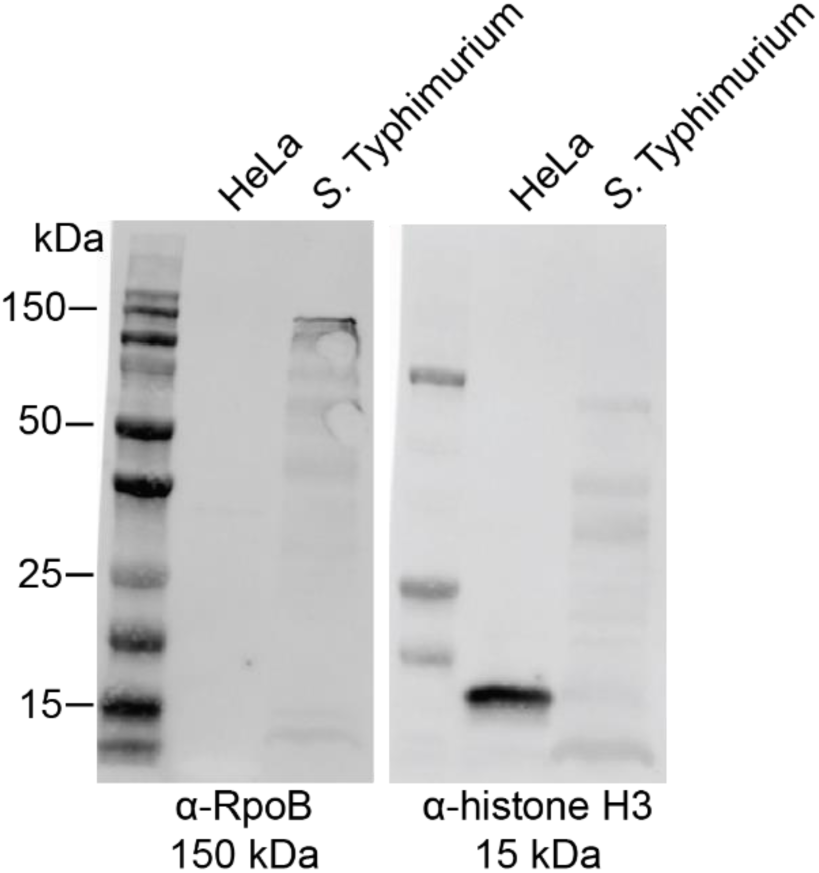
RpoB antibody is specific to bacterial protein. Western blot showing that the native RpoB antibody used in these studies does not cross-react to proteins specific to HeLa cells. HeLa whole cell lysate was prepared in RIPA buffer and diluted into Laemmli sample buffer. *S.* Typhimurium whole cell lysate was prepared in lysis buffer (10 mM Tris-HCl pH = 7, 10 mM EDTA, 1X protease inhibitor, 0.1 mg/mL lysozyme) and diluted into Laemmli sample buffer. Lysates were boiled for 10 min and loaded onto a 12% Mini-PROTEAN TGX gel (Bio-Rad). Proteins were subsequently transferred to a PVDF nitrocellulose membrane (Bio-Rad). The membrane was blocked for 1 h at room temperature with Intercept PBS blocking buffer (LI-COR) before being immunoblotted with anti-RpoB (8RB13, Thermo) and anti-Histone H3 (PA5-16183, Thermo) antibodies overnight at 4°C. The blot was inclubated with secondary antibodies IRDye-680RD and 800CW (LI-COR) for 30 min at room temperature before being imaged on an Odyssey Imager (LI-COR).

**S2 Figure.**
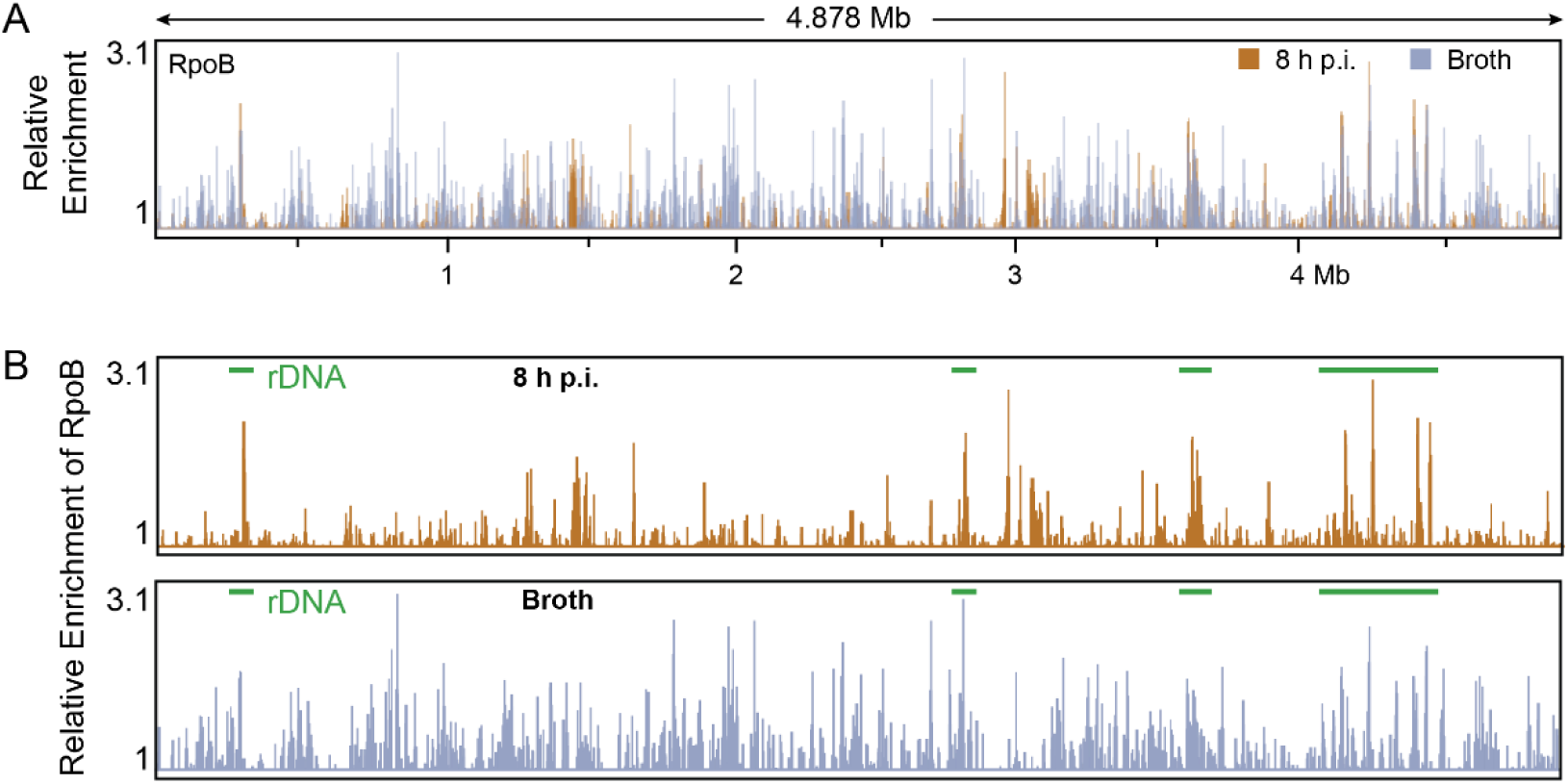
RpoB ChIP-seq signal from broth and infection, related to Figure 2. (A) Representative overlay of *S.* Typhimurium RNAP occupancy as determined by ChIP-seq of RpoB (the beta subunit of RNAP) 8 h post infection (p.i.) of HeLa cells or from cells grown in broth culture. (B) Reads mapped to the *Salmonella* genome display areas of high RNAP enrichment at regions of rDNA in both conditions (green bars). Relative enrichment is defined as the ratio of IP and total read counts.

**S1 Table.** RNAP is significantly enriched at virulence genes during infection. Loci where RpoB was significantly enriched at least two-fold in *S.* Typhimurium 8 h post infection as determined by ChIP-seq. Each value represents the average fold enrichment (‘signalValue’ output of macs2) for at least two biological replicates.

| Locus | Fold Enrichment | Localization | Function | Reference |
| --- | --- | --- | --- | --- |
| *mig-14* | 2.82 | Chromosome | antimicrobial peptide resistance | 1 |
| *virK* | 2.79 | Chromosome | antimicrobial peptide resistance | 2 |
| *sifB* | 2.70 | Chromosome | putatively contributes to the formation of the intracellular replication niche | 3 |
| *sitD* | 2.62 | SPI-1 | resistance to nitrosative stress | 4 |
| *SL1344_1344* | 2.57 | Chromosome | putative component of the type III secretion system (T3SS-2) apparatus | 5 |
| *ssaK* | 2.57 | SPI-2 | type III secretion system (T3SS-2) apparatus | 6 |
| *sitC* | 2.53 | SPI-1 | resistance to nitrosative stress | 4 |
| *pdgL* | 2.53 | Chromosome | putative role in cell wall/membrane biogenesis | 7 |
| *SL1344_2902* | 2.49 | Chromosome | hypothetical protein |  |
| *rplC* | 2.49 | Chromosome | ribosomal protein L3 |  |
| *ssaH* | 2.46 | SPI-2 | putative role in type III secretion system apparatus | 8 |
| *ssaI* | 2.46 | SPI-2 | putative role in type III secretion system apparatus | 8 |
| *ssaJ* | 2.46 | SPI-2 | type III secretion system (T3SS-2) apparatus | 6 |
| *pipB2* | 2.41 | Chromosome | kinesin accumulation in the SCV | 9 |
| *ssaE* | 2.40 | SPI-2 | chaperone | 10 |
| *SL1344_1530A* | 2.37 | Chromosome | hypothetical protein |  |
| *ugtL* | 2.37 | Chromosome | PhoP activation | 11 |
| *sitA* | 2.36 | SPI-1 | resistance to nitrosative stress | 4 |
| *sitB* | 2.36 | SPI-1 | resistance to nitrosative stress | 4 |
| *sseB* | 2.34 | SPI-2 | type III secretion system (T3SS-2) translocon | 12 |
| *sscA* | 2.34 | SPI-2 | chaperone | 13 |
| *sseD* | 2.33 | SPI-2 | type III secretion system (T3SS-2) translocon | 12 |
| *sseK/sseK3* | 2.27 | Chromosome | promote intracellular survival | 14 |
| *sseC* | 2.20 | SPI-2 | type III secretion system (T3SS-2) translocon | 12 |
| *ssaL* | 2.19 | SPI-2 | type III secretion system (T3SS-2) apparatus | 6 |
| *mgtC* | 2.18 | SPI-3 | maintenance of ATP homeostasis | 15 |
| *ssaR* | 2.18 | SPI-2 | type III secretion system (T3SS-2) apparatus | 6 |
| *sseG* | 2.17 | SPI-2 | SIF biogenesis and SCV maintenance | 16 |
| *iscS* | 2.15 | Chromosome | tRNA synthesis | 17 |
| *prgJ* | 2.15 | SPI-1 | type III secretion system (T3SS-1) | 18 |
| *prgI* | 2.15 | SPI-1 | type III secretion system (T3SS-1) | 18 |
| *ssaQ* | 2.10 | SPI-2 | type III secretion system (T3SS-2) apparatus | 19 |

---

**S2 Table.** RNAP is significantly enriched at some of the same genes in broth and infection. Loci where RpoB was significantly enriched at least two-fold in *S.* Typhimurium in cells grown in broth and 8 h post infection, as determined by ChIP-seq. Each value represents the average fold enrichment (‘signalValue’ output of macs2) for at least two biological replicates.

| Locus | Fold Enrichment |  |
| --- | --- | --- |
|  | Broth | Infection |
| *yiaG* | 2.01 | 3.44 |
| *cspA* | 2.01 | 3.44 |
| *cueP* | 2.01 | 3.44 |
| *deaD* | 2.47 | 2.93 |
| *yrbN* | 2.47 | 2.93 |
| *nlpI* | 2.47 | 2.93 |
| *pnp* | 2.47 | 2.90 |
| *cspE (cspC?)* | 2.23 | 2.84 |
| *yobF* | 2.23 | 2.84 |
| *SL1344_RS09185* | 2.23 | 2.82 |
| *mgrB (yobG)* | 2.23 | 2.82 |
| *yqcC* | 2.75 | 2.69 |
| *hdfR* | 2.33 | 2.50 |
| *trxA* | 2.07 | 2.35 |
| *rhoL* | 2.07 | 2.35 |
| *rho* | 2.07 | 2.35 |
| *rpsD* | 2.08 | 2.33 |
| *rpsK* | 2.08 | 2.33 |
| *rpsM* | 2.08 | 2.33 |
| *rpmJ* | 2.08 | 2.33 |
| *rplN* | 2.08 | 2.31 |
| *rplO* | 2.08 | 2.27 |
| *secY/prlA* | 2.08 | 2.26 |
| *rpoA/pez* | 2.08 | 2.18 |
| *rpsC* | 2.21 | 2.18 |
| *rplV* | 2.21 | 2.18 |
| *rplB* | 2.21 | 2.13 |
| *rpoH* | 2.07 | 2.12 |
| *rluD* | 2.37 | 2.08 |
| *pgeF* | 2.21 | 2.03 |

**S3 Figure.**
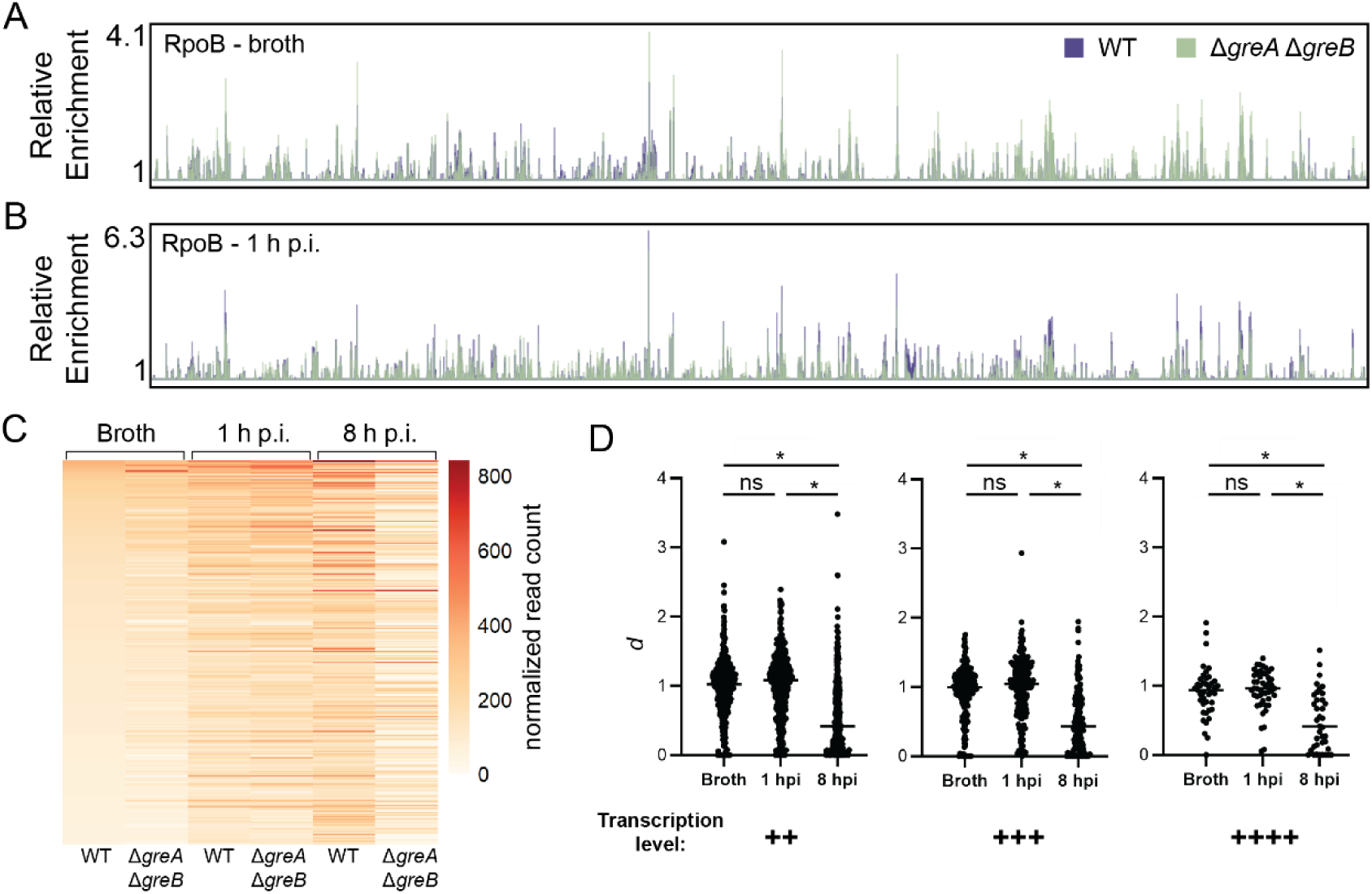
RNAP occupancy changes in the presence and absence of Gre factors, related to Figure 3. Representative RNAP occupancy profile for wild-type (WT) cells or cells lacking Gre factors (Δ*greA* Δ*greB*) in cells (A) grown in broth and (B) at 1 h post infection (h p.i.) as determined by ChIP-seq of RpoB. Relative enrichment is defined as the ratio of IP and total read counts. (C) RNAP occupancy changes visualized as a heat map, where every horizontal line represents the normalized read count for the same gene across each condition. Only top-transcribed genes that arise in all three conditions are plotted (283 genes). Each value represented is the average of three biological replicates. (D) Quantification of *d* ratios calculated for each gene falling within each transcription level (see S3 Table). The number of genes belonging to each condition within each transcription level is as follows: ++, 354 (broth), 412 (1 h p.i.), 405 (8 h p.i.); +++, 160 (broth), 164 (1 h p.i.), 147 (8 h p.i.); +++, 37 (broth), 42 (1 h p.i.), 38 (8 h p.i.). Each value represents the average of three biological replicates. \**p*<0.0001, one-way ANOVA.

*Excel file*

**S3 Table. Top transcribed genes in cells grown in broth, 1 h p.i., and 8 h p.i.** List of the top transcribed genes as determined by k-means clustering for each condition. Values represent the average normalized read count and the ratio of the normalized read counts of three biological replicates. This table also lists the 283 top transcribed genes that arise in all three conditions.

**S4 Figure.**
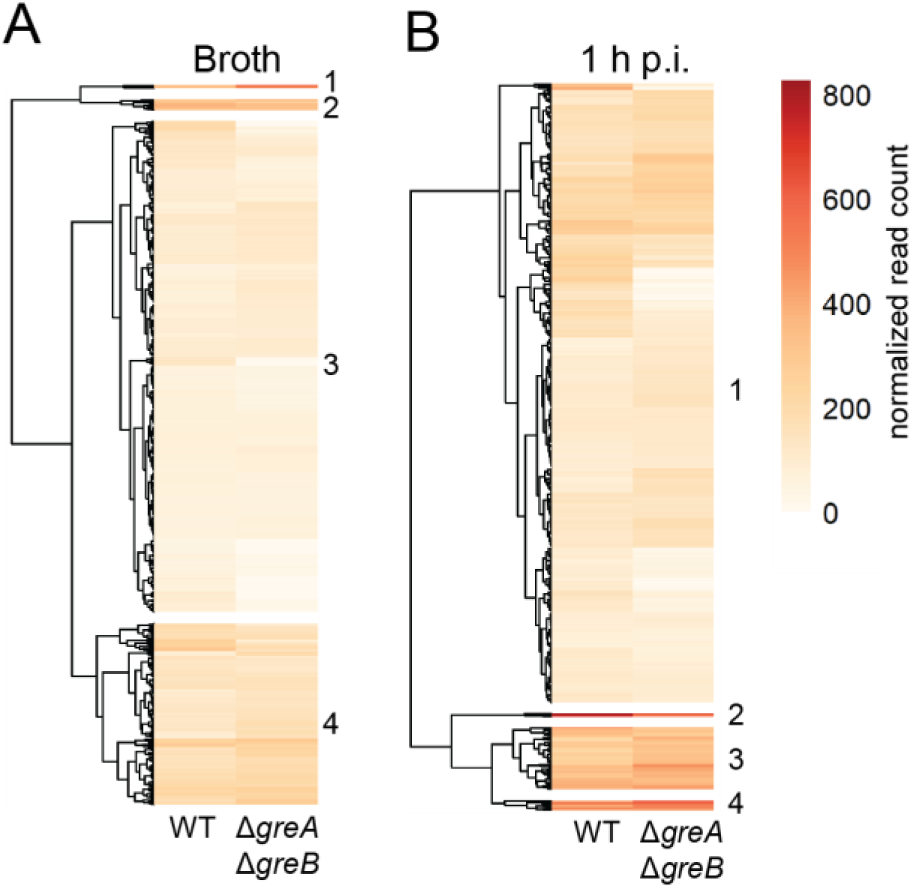
RNAP occupancy differences in broth and 1 h p.i., related to Figure 4. RNAP occupancy changes in cells grown in broth and at 1 h post infection (h p.i.) in the absence of Gre factors visualized as a heat map. Each value represents the average of three biological replicates. Hierarchical clustering (numbered 1-4) was performed using the pheatmap function in RStudio (see Table S4).

*Excel file*

**S4 Table. RNAP occupancy changes for the top transcribed genes as categorized by hierarchical clustering.** Lists the top transcribed genes for each condition that call into each hierarchical cluster, as determined by pheatmap library in R. Values represent the average normalized read count and the ratio of the normalized read counts of three biological replicates. Also lists the manually annotated functions of genes in clusters three and four for the 8 h post infection (p.i.) condition.

**S5 Figure.**
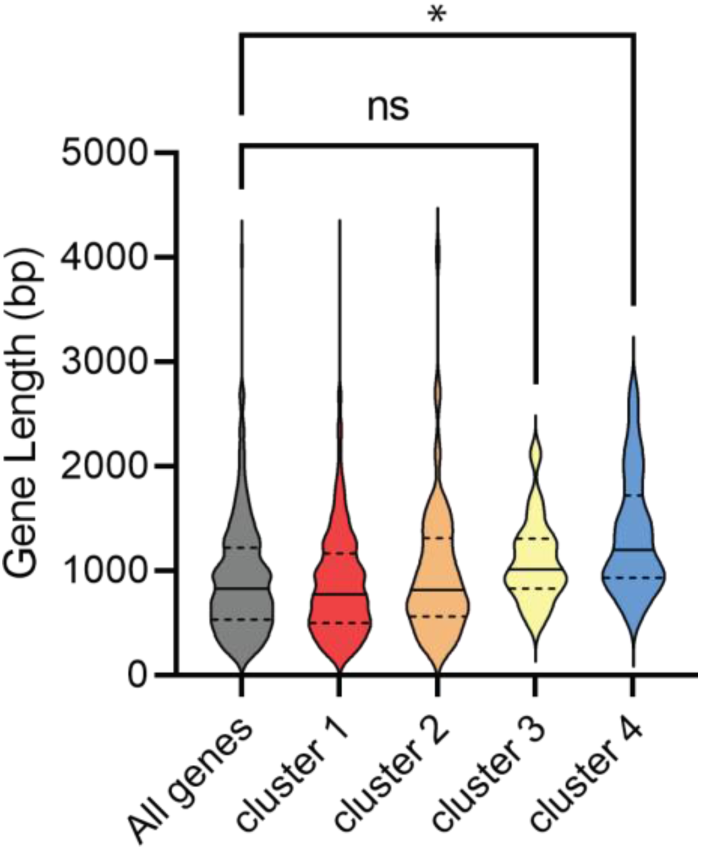
Longer genes experience more prevalent backtracking, related to S4 Table. All top transcribed genes at 8 h post infection were sorted by hierarchical clustering using the pheatmap function in RStudio (590 genes total). The average gene length was calculated for genes in each cluster. \**p*<0.001, ns: not significant, one-way ANOVA.

**S6 Figure.**
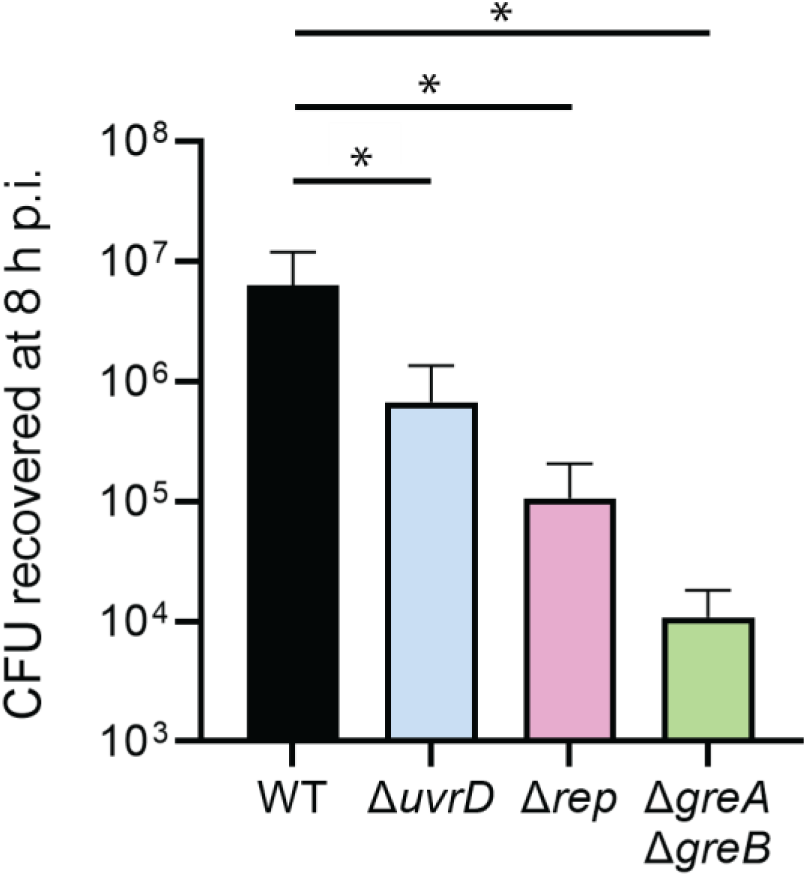
Cells lacking key conflict resolution factors exhibit significant growth defects during infection. HeLa cells were infected with *S.* Typhimurium cells lacking the indicated genes. Bacteria were harvested at 8 h post infection (p.i.) and plated for CFU enumeration. \**p*<0.05, one-way ANOVA.

**S5 Table.**
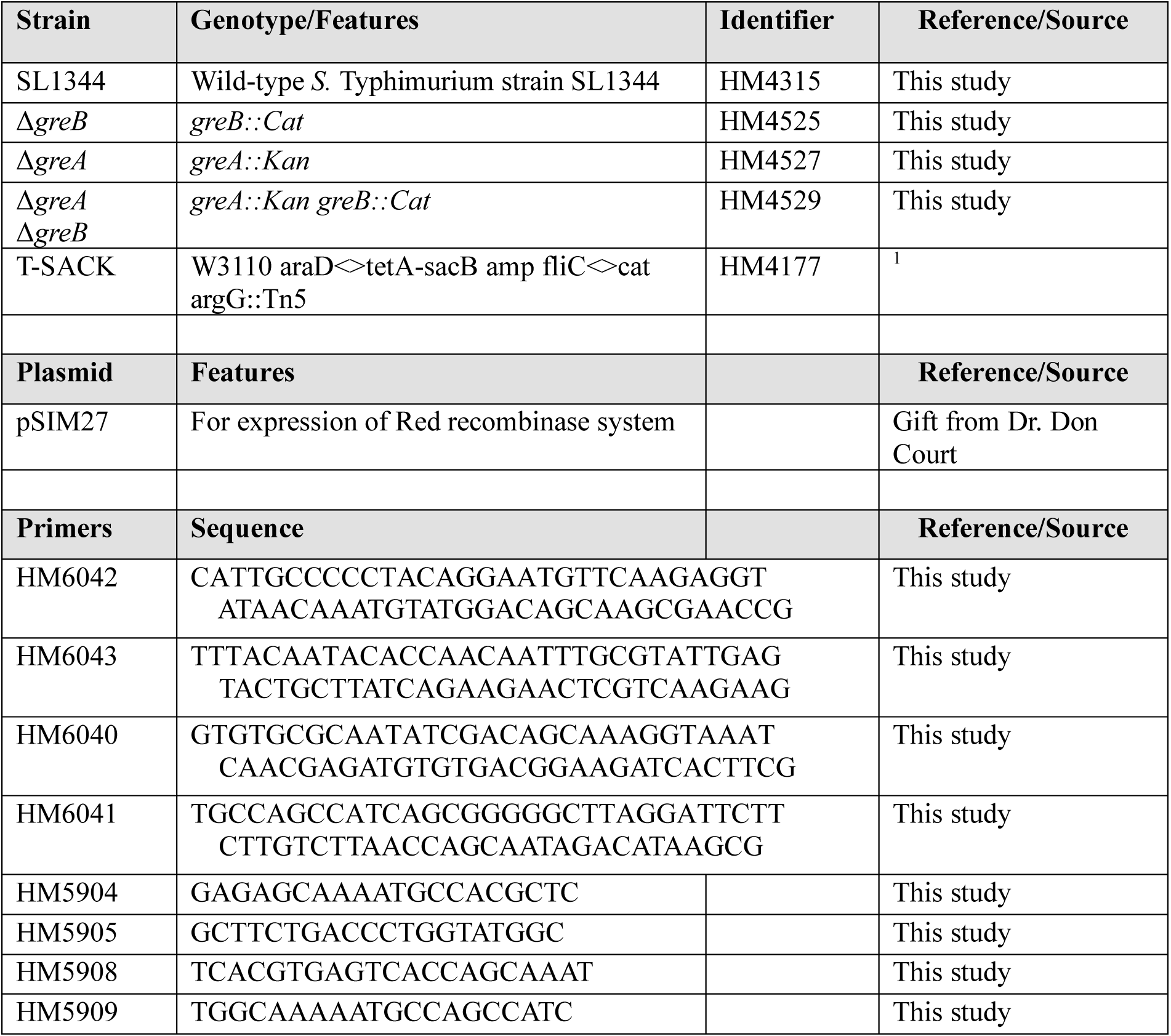
Strains, plasmids, and primers used in this study.

